# Thermoplasmonic modulation of cardiomyocytes activity with local temperature read-out

**DOI:** 10.1101/2023.08.21.554091

**Authors:** Fanglei Guo, Wei Yu, Olivier Deschaume, Stijn Jooken, Liwang Liu, Qi Wei, Wim Thielemans, Carmen Bartic

## Abstract

Cardiomyocyte beating rate modulation is required in multiple *in vitro* assays and it is usually done by electrical or optogenetic methods. In this work, we develop and characterize a light-based strategy to modulate the cardiomyocyte beating rate by near-infrared (NIR) controlled plasmonic stimulation. For this purpose, gold nanorods (GNRs) acting as plasmonic heaters and silica-coated quantum dots (QDs) as nanothermometers are attached to surfaces used for cardiomyocyte cultures, while the cell electrophysiological activity is visualized by monitoring calcium transients via calcium imaging. This system is capable of modulating cardiac activity with near infrared-controllable plasmonic heating while measuring local temperature changes owing to the temperature-dependent fluorescence of silica-coated QDs.

**Table of Contents (TOC):** 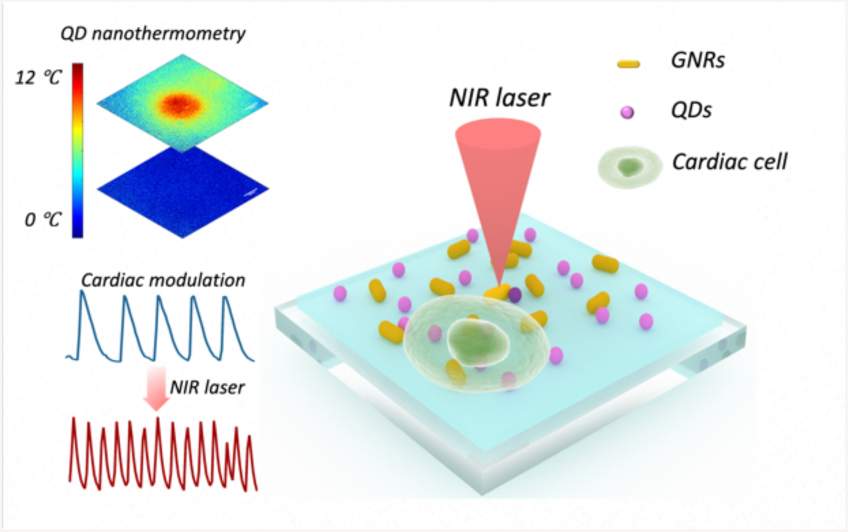

## 1. INTRODUCTION

Modulating cardiomyocyte beating rate is essential in various *in vitro* functional assays used for instance in pharmacological compound screening and cardiac toxicity assays.^1^ Electrical-^2–4^ and optogenetic^5–10^ activity modulations are widely used strategies for cardiac activity control. However, electrical modulation protocols are either invasive or have low spatial resolution, while optogenetic stimulation requires genetic manipulations. Modulating cardiac function with light using photoswitchable compounds without genetic manipulation is also an emerging approach in cardiac photopharmacology studies, where small molecules with photochromic properties are utilized to control cardiomyocyte function by regulating G protein-coupled receptors (GPCR) activity.^11,12^ Apart from these, non-optogenetic photomodulation based on external photo-responsive materials is another promising alternative for cardiac activity modulation, as it can regulate the activity of cells and tissues non-invasively and with a high spatial resolution, even in 3D cell cultures. Various materials, such as silicon nanowires,^13^ conjugated polymer films,^14^ graphene,^15^ porous metamaterials,^16^ gold nanoshells (GNSs)^17^ and gold nanorods (GNRs)^18–20^ have been proposed for optical muscle and cardiac activity modulation based on either photoelectric or photothermal effects. GNRs are good candidates for photothermal cell activity modulation, owing to their thermoplasmonic properties, tunable surface functionality, good biocompatibility, and facile attachment to various cell scaffold materials.^21^ The longitudinal localized surface plasmon resonance (LSPR) band of GNRs can be tuned over a large wavelength range from the visible to the near-infrared (NIR) and infrared (IR) regions, through proper regulation of the rod aspect ratio.^22^ Previous studies have also shown that by adjusting the GNR size and excitation power, the local temperature increment around these particles can range from a few degrees to a few hundred degrees.^23^ As a proof of concept, Gentemann *et al.* treated HL-1 cultures with unmodified 200 nm gold spheres and showed that high power irradiation with a 532 nm picosecond pulsed light source could induce calcium shock in directly irradiated cells and calcium oscillations in other cells in the culture.^18^ Marino *et al.* exploited GNSs with NIR to generate mild intracellular temperature changes in muscle cells, resulting in myotube contraction modulations without calcium fluxes involved.^17^ Recently, Fang *et al.* proposed using photothermal stimulation of plasmonic nanostructured electrode arrays to restore the cardiomyocyte beat rate in a bradyarrhythmia in vitro model.^19^ These studies demonstrated for the first time the possibility of using plasmonic stimulation as a cardiac modulation method via non-invasive optical stimulation.

However, various studies report contradictory effects of plasmonic stimulation on cell activities – with several reporting inhibition of electrical activity of electrogenic cells^24,25^ and calcium transients independent modulation^17^ under NIR illumination. There is thus a lack of understanding of specific modulation effects, since LSPR can not only lead to heat dissipation, but could in principle also induce electrical modulation through photocurrent generation, as proposed by Bruno *et al*.^16^. In order to understand the role of these effects, precise readout of local temperature changes is essential.

In this paper, we report on a platform where NIR-controlled plasmonic stimulation of cardiomyocytes is performed while monitoring the resulting local extracellular temperature by quantum dot (QD) nanothermometers. Accurate monitoring of the extracellular temperature during thermoplasmonic modulation allows for resolving the role of temperature in activity modulation and precisely adjusting the stimulation protocols to control cell activity.

QDs are interesting for nanothermometry due to their unique, temperature-dependent, photoluminescence properties.^26–28^ Compared to conventional organic dyes that bleach quickly under constant illumination, QDs have a higher photostability and absorption cross-sections, retaining high emission intensities after extended illumination periods in biological systems.^29^

Specifically, we developed and calibrated GNR and QD-decorated substrates that enable modulating cardiomyocyte activity while measuring local extracellular temperature changes with single-cell resolution. Surface immobilized nanoparticles prevent the cellular uptake of NPs, which usually occurs when using NP dispersions and in this way the NP localization in respect to temperature sensitive cellular structures is well defined. In this all-optical platform, we utilized calcium imaging to monitor calcium transients as an indirect indication of cardiomyocyte beating activity. We calibrated first the temperature-induced activity changes by gradually modifying the culture temperature in a thermostated bath. Afterwards, we applied NIR laser irradiation, and by nanothermometry, we calibrated the local temperature changes that can go up to 10°C for moderate irradiance levels. Our experiments demonstrate that NIR plasmonic stimulation can reversibly modulate HL-1 cardiomyocyte activity, and that the activity modulation is due to photothermal effects. Since such light-addressable nanoparticle structures are constructed by successive self-assembly steps, they can be easily integrated into 3D cell matrices to achieve photonic cellular activity modulation coupled with local temperature readout in cardiac tissue models.

## 2. EXPERIMENTAL SECTION

### Materials

All materials were used as received without any further purification. Cetyltrimethylammonium bromide (CTAB), polystyrene sulfonate (PSS), Poly-L-lysine hydrobromide (PLL), 1-ethyl-3-(3-dimethylaminopropyl)-carbodiimide (EDC), sulfo-N-hydroxysulfosuccinimide (NHS), Pluronic F-127, dimethyl sulfoxide (DMSO), HL-1 cardiac muscle cell line, Fetal Bovine Serum (FBS), Triton X-100 and Norepinephrine were purchased from Merck. Tri-sodium citrate was obtained from Acros Organics, (3-Aminopropyl)triethoxysilane (APTES) was obtained from ABCR, 4-(2-hydroxyethyl)piperazine-1-ethanesulfonic acid (HEPES), penicillin/streptomycin, Glutamax, Fluo-3 AM, Dulbecco’s Phosphate Buffered Saline (DPBS), Horse serum, Connexin 43 primary Antibody (Connexin43 (1A)), Goat anti-Mouse IgG (H+L) Cross-Adsorbed Secondary Antibody, Alexa Fluor™ 647 and Hoechst were purchased from Fisher Scientific. Sulfuric acid (97%) and hydrogen peroxide (31%) were from Chem Lab. Ultrapure (UP) water (>18.2 MΩ) was used for all experiments and produced with a Sartorius Stedim Arium Pro VF system.

### Cleaning of Substrates

All silicon and glass substrates used in this study were first freshly cleaned by immersion for 30 min in piranha solution (H_2_SO_4_ (98%)/H_2_O_2_ (30%) 3/1 v/v), rinsed extensively in ultrapure water, and dried under a nitrogen flow before further use.

### Gold Nanorods Synthesis

Gold nanorods (GNRs) were synthesized following a seed-mediated protocol.^30,31^ The detailed synthesis protocol is described in our previous work.^32^ After synthesis, GNRs were purified by three centrifugation steps using an Eppendorf minispin centrifuge (30 min at 8000 rpm), replacing each time the supernatant with 5 × 10^−3^ M CTAB. To further improve the GNR biocompatibility, the GNRs were then transferred to a 5 × 10^−3^ M sodium citrate solution following a capping agent exchange protocol.^33^ Briefly, 2mL CTAB-stabilized GNRs were first centrifuged four times at 8000 rpm for 30 minutes, re-suspending the pellet in 1.5 mg mL^−1^ polystyrene sulfonate (PSS) solution and aging for 1 hour for the first two rounds, and re-suspending the pellet in 5 mM sodium citrate, before aging for at least 12 hours for the last two. The resulting citrate-stabilized suspension was then stored until further use. Citrate stabilized GNRs (cit-GNRs) were characterized with atomic force microscopy (AFM) and UV-visible spectroscopy to determine their morphology and spectral properties (Figure S1, Supporting Information). From AFM images in S1a, cit-GNRs have a length of 55.4 ± 12.9 nm and a width of 16.9 ± 1.9 nm, with a longitudinal plasmon absorption band (Figure S1b, Supporting Information) at 750 nm.

### CdSe/CdS@SiO_2_ QDs synthesis and water solubilization

CdSe/CdS@SiO_2_ QDs were first synthesized through the “flash” method from Aubert et al.,^34^ followed with a carboxylated silica encapsulation procedure^35^ and PEG termination (9-12 PE units),^36,37^ the core diameter of synthesized CdSe/CdS@SiO_2_ QDs is 9 nm, to which a 8.5 nm thick SiO_2_ shell is added, as demonstrated by TEM. Absoption and emission spectra, together with a TEM image, are shown in Figure S2, Supporting Information.

### GNR-QD-modified substrates preparation

Glass coverslips were first cut into 1cm wide squares to fit in the temperature-controlled imaging chamber (JPK Biocell^TM^, Bruker, Germany), then treated with piranha solution for 30 min, extensively rinsed with ultrapure water, and dried with nitrogen gas. Next, cleaned coverslips were silanized with APTES.^38^ In brief, the coverslips were immersed in ethanol for 5 minutes and dried with nitrogen gas, then immersed in a 5% (v/v) APTES solution in ethanol for 20 min, rinsed with ethanol, and finally dried. The silanized glass coverslips were then immersed in a 6% (v/v) aqueous acetic acid solution for 60 min, rinsed with ultrapure water, and dried with nitrogen gas. After 24-48 hours of storage at 10 °C, the coverslips were ready for GNR and QD modification. GNRs were attached to the surface through electrostatic absorption. For this purpose, the APTES-modified coverslips were immersed in 5.8 × 10^−10^ M cit-GNR solution overnight, rinsed with ultrapure water, and dried with nitrogen gas (AFM images shown in Figure S3, Supporting Information). CdSe/CdS@SiO_2_ QDs were then covalently bound to APTES modified coverslips through EDC/NHS reaction. Briefly, 100 μL of 1 × 10^−7^M CdSe/CdS@SiO_2_ QDs solution in water was first mixed with 900 μL of 1 × 10^−1^ M, pH 6.5 MES buffer containing 5 × 10^−3^ M EDC and 5 × 10^−3^ M NHS. Modified coverslips were immersed in the EDC/NHS solution for at least 4 hours, rinsed twice with ultrapure water, and dried with nitrogen gas. The prepared samples were then ready for the HL-1 cell culture.

### HL-1 cell culture

HL-1 cells were purchased from Merck and cultured according to the standard cell culture protocol provided by the supplier. Cells were cultured in Claycomb medium supplemented with 10% FBS, 100 μg penicillin/streptomycin, 1 × 10^−4^ M Norepinephrine, and 2 × 10^−3^ M Glutamax in a Binder incubator at 37 °C and 5 % CO_2_. The T25 culture flasks were precoated using 1 mL of 0.02% gelatin and 0.005 mg mL^−1^ fibronectin in water for at least 1h at 37 °C, followed by a rinse with DPBS. The cells were passaged 1:2 two times a week upon confluency. Following a rinse with DPBS without Ca^2+^ and Mg^2+^, the cells were dissociated through the addition of 1 mL of 0.05 % trypsin/EDTA for 1 min at 37 °C, which was then replaced by 1 mL of 0.05 % trypsin/EDTA for 2 more minutes at 37 °C. Then 1mL of soybean trypsin inhibitor was added to quench the enzymatic reaction. Cells were then centrifuged for 5 min at 500×g and resuspended 1:2 in 5 mL of supplemented Claycomb medium in a gelatin/fibronectin-coated T25 flask.

For microscopy, piranha-cleaned glass coverslips were first cut into 1 cm wide squares, immersed in 70 % (v/v) ethanol for 10 min for sterilization, placed in a 24 well plate, and seeded with either 10000/cm^2^ or 100000/cm^2^ HL-1 cells in 2 mL growth medium. The culture medium was refreshed every day and after 2 days the samples were used for experimentation.

For the experiments involving plasmonic stimulation, cell adhesion on the GNR-QD modified surfaces was improved by coating with 1 mg/mL PLL in 1 × 10^−1^ M, pH 8.5 borate buffer for 10 min, rinsed with ultrapure water, and dried with nitrogen gas. Before seeding with the HL-1 cells, Modified surfaces were sterilized by immersion in 70 % (v/v) ethanol, followed by a rinse with DPBS. Then, 10000/cm^2^, 50000/cm^2^, or 100000/cm^2^ HL-1 cells were seeded on the sample surface in a 24-well plate and maintained with 2 mL Claycomb medium. The different initial seeding densities correspond to the ‘single cells’, ‘small clusters’ and ‘confluent cells’ samples discussed in the article. The culture medium was refreshed every day and after 2 days the samples were used for measurements.

### Calcium imaging

HL-1 activity was studied through calcium imaging, which was performed using the calcium indicator Fluo-3 AM according to the manufacturer’s instructions.^39^ 50 μG Fluo-3 solution was dissolved in 14 μL DMSO containing 20 % (w/w) Pluronic F-127 and diluted 1000 times with HEPES buffer (where mentioned in the text, HEPES always refers to 1 × 10^−2^ M, pH 7.4 HEPES buffer supplemented with 1.4 × 10^−1^ M NaCl, 5 × 10^−3^ M KCl, 2 × 10^−3^ M CaCl_2_ and 1 × 10^−3^ M MgCl_2_, osmolarity 292 mOsm/L). Samples were incubated in Fluo-3 solution for 30 min, rinsed twice with HEPES, and left for an additional 30 min in HEPES before performing microscopy.

### Fluorescence microscopy & NIR-induced plasmonic heating

For global sample heating, Fluo-3 AM-loaded HL-1 cells cultured on clean glass coverslips were placed in the JPK Biocell^TM^ temperature-controlled imaging chamber in HEPES buffer. The chamber temperature was set to 21, 26, 30, 35, 40, and 45 °C to record temperature-dependent calcium transients. The Fluo-3 emission was recorded with an inverted fluorescence microscope (Olympus IX81) using a U-MWIB3 Olympus filter cube, containing a 460-495 nm bandpass excitation filter, a 520-550 nm bandpass emission filter, and a 505 nm dichroic mirror. A 20x objective lens (Olympus LUCPLFLN20XRC, NA 0.45) was applied for fluorescence collection and a Hamamatsu ORCA-Flash 4.0 v2 scientific camera was used for image acquisition with an exposure time of 200 ms. Calcium signal displayed from indicated area with averaged fluorescence intensity was aquired by ImageJ.^40^

To calibrate the temperature sensitivity of CdSe/CdS@SiO_2_ QDs on 2D surfaces, the same set-up was used as for Fluo-3 monitoring, but with a U-MWIG3 Olympus filter cube, which contains a 530-550 nm bandpass excitation filter, a 575 nm long pass emission filter, and a dichromatic mirror (short wavelength reflection) with a cut-off wavelength of 570 nm.

For the calibration of CdSe/CdS@SiO_2_ QDs in solution, CdSe/CdS@SiO2 QDs were dispersed in 1 × 10^−2^ M, pH 7.4 HEPES buffer and sealed in a quartz cuvette. The cuvette was mounted in an optical cryostat (Optistat-DN-V, Oxford Instruments) to control the temperature. A CW-532 nm probe laser (06-DPL, Cobolt) was used to excite the QDs and a fiber-coupled spectrometer (USB 4000, Ocean Optics) was used to collect the emission.

For NIR-induced GNR heating of HL-1 cells, a 785 nm collimated NIR laser was added to the aforementioned setup. The NIR laser was fiber-coupled to a 10x objective lens (Olympus UPLANFL10X, NA 0.3) and mounted in the ‘upright’ position above the sample stage as shown in Figure 1.

**Figure 1.**
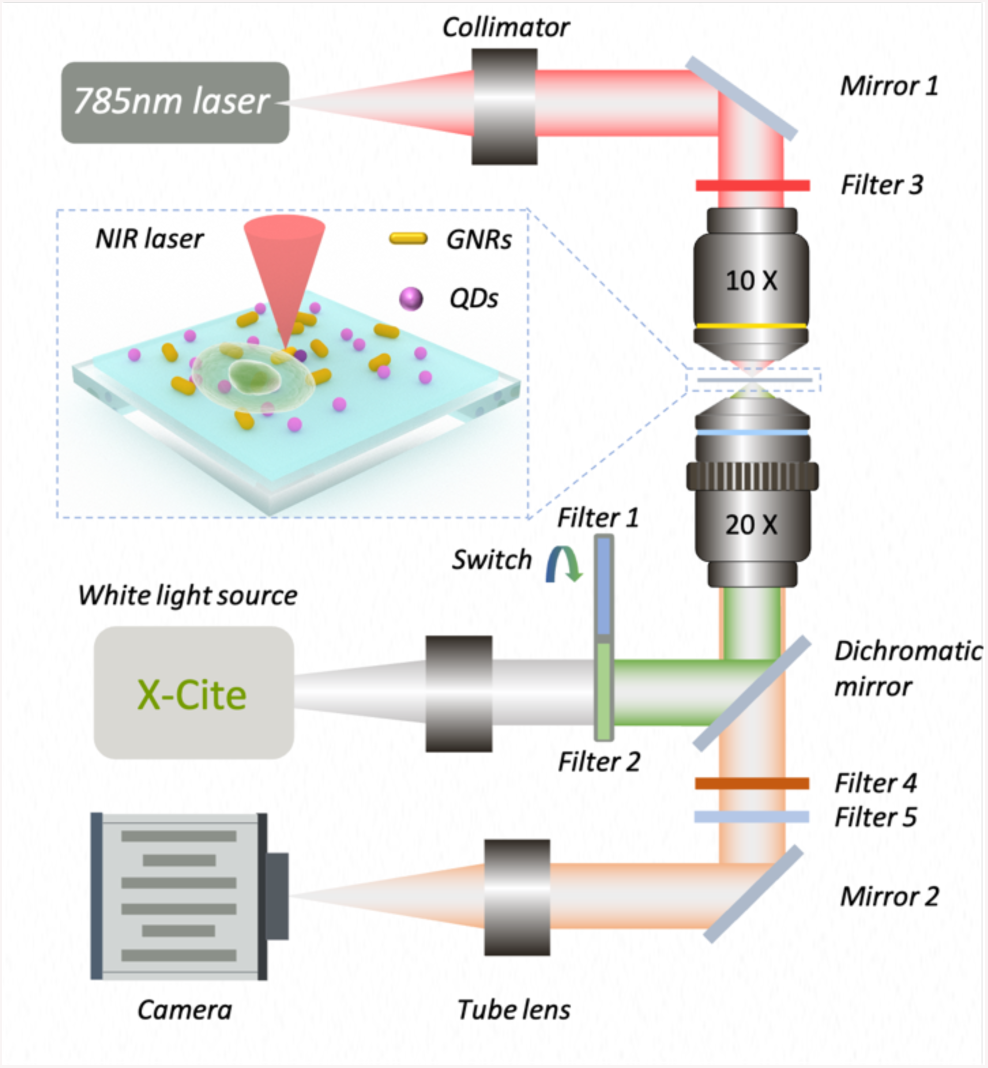
Scheme of the set-up used for cell activity modulation and temperature read-out.

### Immunofluorescence staining and confocal microscopy

Immunoconfocal microscopy was used to observe the formation of gap-junction protein Connexin 43 in 2 DIV cultured HL-1 single-cell samples and small cluster samples. Cells cultured on clean glass were fixated with 4% PFA at room temperature for 10 minutes and washed three times with DPBS. Following fixation, samples were incubated in 0.3% (v/v) Triton X-100 DPBS for 10 minutes at room temperature, followed by rinsing three times with DPBS. Afterward, samples were incubated with 10% (v/v) horse serum at room temperature for 1 hour to block unspecific binding sites followed by labeling with 5 μg mL^−1^ primary antibody at 4 °C overnight. After rinsing five times with DPBS, samples were incubated with 10 μg mL^−1^ fluorescent dye-conjugated secondary antibody and 200 μM Hoechst solution at room temperature for 90 minutes, followed by five rinsing steps with DPBS. Confocal microscopy of immunolabelled HL-1 culture was carried out using a Leica SP8 dive microscope with a 25x water immersion objective lens (HC FLUOTAR L 25x/0.95 W VISIR).

### Statistical analysis

Paired t-test and Two-way ANOVA were analyzed with GraphPad Prism, and the p-values calculated from paired t-test were indicated as follow, (*) p < 0.05, (**) p < 0.01, (***) p < 0.001, (****) p < 0.0001, (ns) not significant.GNR produced temperature maps were calculated and generated with a MATLAB script as discussed in Section 3.4, second paragraph.

## 3. RESULTS AND DISCUSSION

### 3.1 Thermoplasmonic modulation and local temperature read-out system

To modulate cell activity while monitoring the temperature changes, the nanoparticle-modified substrate was probed using the optical set-up depicted in Figure 1, detailed description is given in Experimental Section. The nano-functionalized substrate was prepared as follows: An APTES-modified glass slide was first coated with 16.9 nm wide and 55.4 nm long citrate-stabilized GNRs (AFM images and UV-visible spectrum shown in Figure S1, Supporting Information) via electrostatic absorption and then silica-coated QDs functionalized with PEG-silane were grafted via N-ethyl-N′-(3-(dimethylamino)propyl) carbodiimide/N-hydroxysuccinimide (EDC/NHS) chemistry (detailed protocol in the Experimental Section). To improve cell adhesion on the surface, a thin layer of poly-L-lysine (PLL) was deposited on the GNR-QD modified substrate before seeding the HL-1 cardiomyocytes. During measurements, the substrate was placed on an inverted fluorescence microscope and NIR stimulation was applied from the top with a collimated laser placed above the sample stage. Two interchangeable filter cubes with green and blue excitation light were employed for QDs and Fluo-3 AM calcium dye-loaded cell observation, respectively.

To study the effect of temperature on the cardiomyocyte beat rate, first, the temperature of the entire culture was varied in a temperature-controlled chamber, and in a second series of experiments, through localized plasmonic heating.

### 3.2 Temperature effect on HL-1 activity depends on the cell density: calibration studies by modifying the global culture temperature

First, the temperature effect on calcium transients was optically monitored in HL-1 cell cultures on clean glass substrates, for different cell densities. For this purpose, the temperature of the culture was set to 5 different values, in a temperature-controlled chamber (JPK biocell^TM^). The cell coverage was controlled by varying the initial number of cells seeded on the substrate. Low cell density samples, i.e., consisting of mainly isolated cells (without connections with others in the culture, Figure 2b, top right image), are further referred to as ‘single cells’ samples, whereas substrates, where cells have reached confluency and are highly interconnected, are referred to as ‘confluent cells’ samples (Figure 2b, bottom right image). Figure 2b shows an example of calcium signals from 2 types of samples cultured at 30°C and their corresponding fluorescence images.

**Figure 2.**
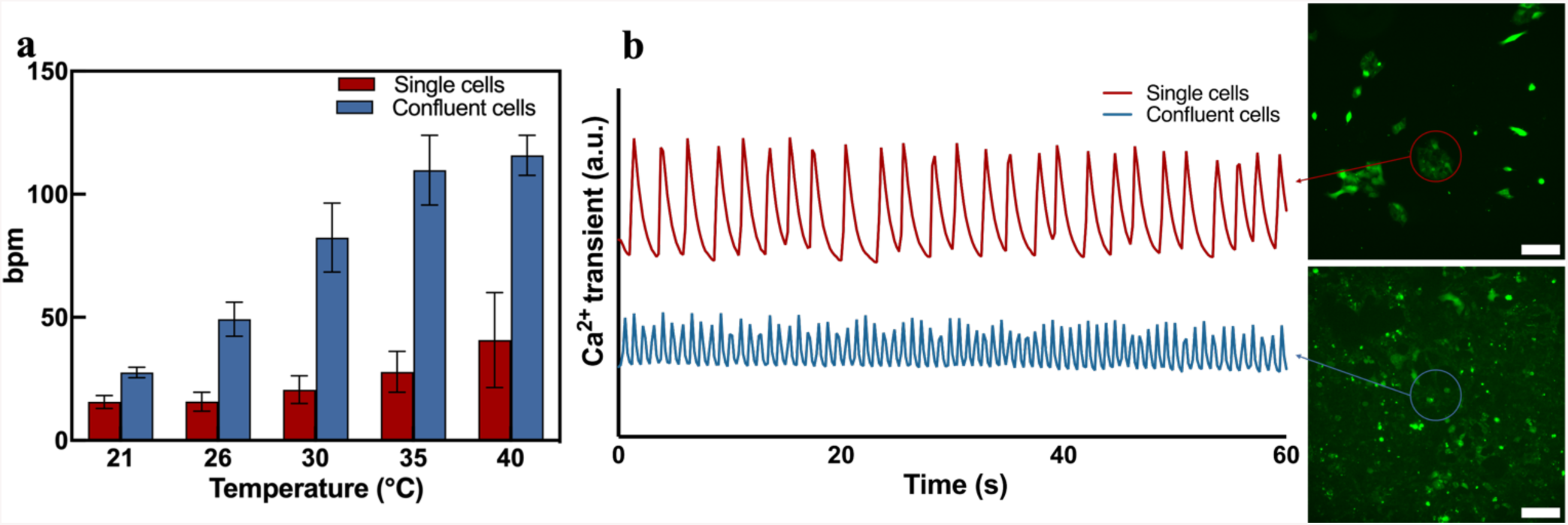
Influence of global temperature (i.e., bath temperature control) on HL-1 beating rate for two culturing conditions: i.e., confluent and low density (i.e., ‘single cell’) cell cultures. **a.** Beating rates of 2 DIV HL-1 cells (2 DIV) as a function of environmental temperatures and cell densities. Two-way ANOVA test: Temperature p < 0.0001, Cell density p < 0.0001. **b.** Calcium transients of ‘single cells’ and ‘confluent cells’ samples at 30°C and the corresponding fluorescence images. Scale bar: 100 µm.

The beating rate of cardiomyocytes increased with the culture chamber temperature, in agreement with previous studies.^41,42^ Although the mechanisms underlying this phenomenon are unclear, they may be primarily related to temperature-dependent ion channel activity. When temperature increases, ion channel activity increase accelerates the Ca^2+^ exchange rate through the cell membrane, resulting in reduced contraction-relaxation sarcomere cycles, and therefore increases the beating rate.^41,43^ Furthermore, from Figure 2a one can see that confluent cells beat faster than isolated cells at all temperatures, and beating rates varied more with temperature changes. The beating rate change with respect to the beating rate at 21°C for confluent samples was also larger than for single cell samples, the largest variation being observed at the highest temperature. At 40°C confluent cells beat approximately 4.2 times faster than when measured at 21 °C, while for isolated cells in low density cultures the beating rate increased about 2.6 times between 21 and 40°C. A two-way ANOVA test was performed on data in Figure 2a taking global temperature and cell density as the two parameters. This gave temperature p < 0.0001 and cell density p < 0.0001, indicating that both global temperature and cell density significantly influence the beating rate. Higher beating rates in cell monolayers may result from confluent cells forming more gap junctions that stabilize and enhance electrical activity of connected cardiomyocytes.^44,45^ Also these experiments show that the temperature sensitivity depends on the interconnectivity level of cardiomyocyte cultures, in agreement with a previous study proposing that immature cardiomyocytes are less sensitive to temperature changes.^41^ Moreover, the temperature-induced change in the beating rate is dependent on the initial beating rate. The activities of fast-beating cells (i.e., in our case cells beating close to 100 bpm) cannot be further increased when the temperature increases beyond 40 °C, while for slower-beating cells, the beating rates keep increasing above 40°C. Finally, this first series of experiments confirm the effect of thermomodulation on cardiomyocyte activity and provide a baseline against which the thermoplasmonic modulation experiments will be compared.

### 3.3 Calibration of Quantum Dot-based local temperature readout

To monitor local temperature changes upon GNR illumination, silica-coated CdSe/CdS@SiO_2_ quantum dots with a core diameter of 9 nm and a 8.5 nm thick SiO_2_ shell were used as thermal nanoprobes (TEM and Extinction/Emission spectra shown in Figure S2, Supporting Information). Prior to these experiments, the QD temperature sensitivity was first calibrated in solution (Figure S4, Supporting Information) as well as after immobilization onto a glass surface (Figure 3), together with the GNRs, to accurately mimic the conditions used for the cell experiments. For both experiments, the QDs were solubilized in the same 1 × 10^−2^ M, pH 7.4 HEPES buffer as used for the HL-1 cell experiments. The QD solutions were illuminated with a 532 nm laser (6.36 mW/mm^2^) and photoluminescence was collected with a spectrometer (as described in the Experimental Section). The QD spectra were recorded at six different temperatures between 30 to 48°C. When increasing the temperature, the QD emission peak showed a gradual red shift and a concomitant decrease in photoluminescence intensity which was (1.76 ± 0.21) % per °C (results shown in Figure S4, Supporting Information).

**Figure 3.**
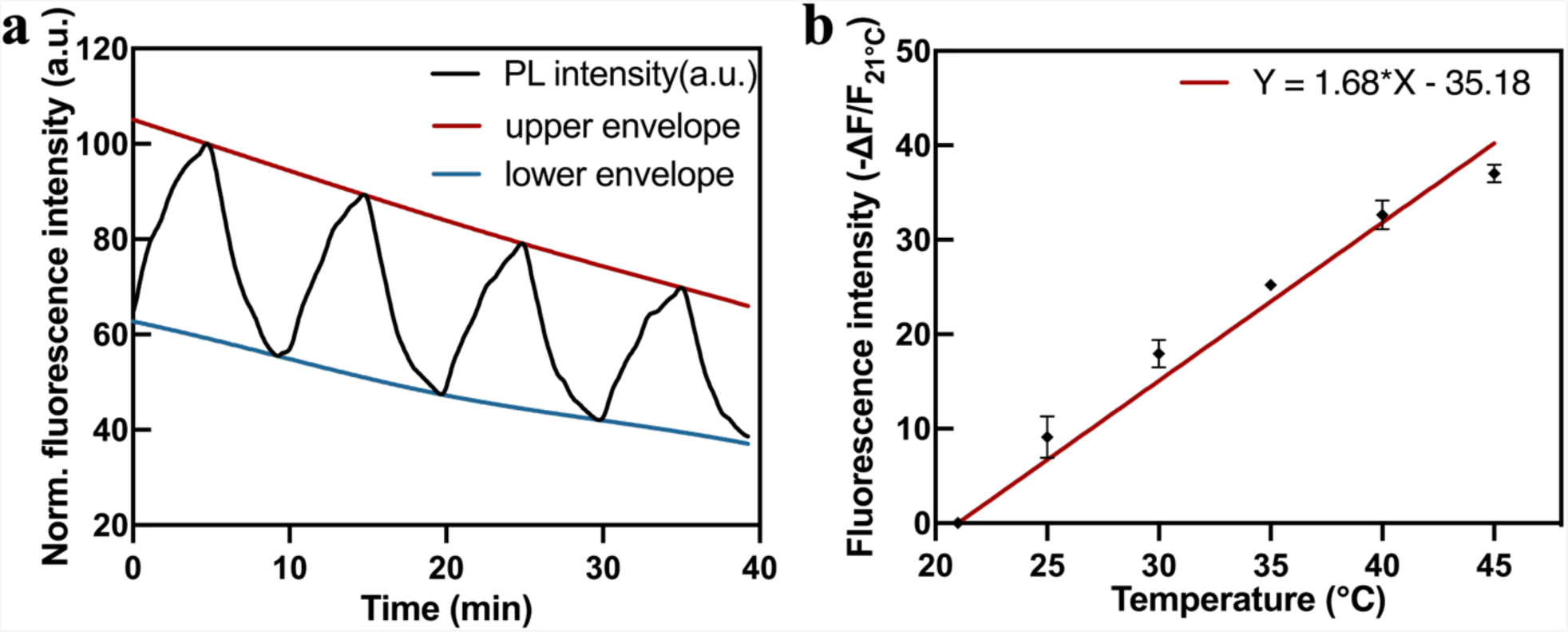
Calibration of CdSe/CdS@SiO_2_ QD temperature sensitivity. **a.** Photoluminescence (PL) intensity profile of GNR-QD mediated substrate in 1 × 10^−2^ M, pH 7.4 HEPES buffer upon temperature oscillation from 21 to 45°C. **b.** Normalized PL instensity change as a function of temperature from 21 to 45°C by 5 °C increments.

Similar experiments were performed with the surface-immobilized QDs sample immersed in 1 × 10^−2^ M, pH 7.4 HEPES buffer. The photoluminescence intensity was acquired from the images recorded with a Hamamatsu Orca Flash CMOS camera while varying the chamber temperature between 21 and 45°C (Figure 3). To study the stability and reversibility of the QD probes, their photoluminescence was recorded by cycling the environmental temperature between 21 and 45 °C (Figure 3a) for four cycles under constant illumination. Under such constant illumination, the QD PL gradually decayed over time, but still remained at about 70% of the initial PL intensity after 40 minutes of constant illumination, without significantly changing the temperature sensitivity. We measured an average photoluminescence intensity change of (42.75 ± 1.18) % for a temperature change of 24°C. In later experiments, the QD thermometers were only illuminated while recording the temperature, typically less than 1 minute, which did not lead to a measurable fluorescence decay. The intensity variation displayed as a function of temperature in Figure 3b was acquired for temperatures varied between 21 °C and 45 °C by 5 °C increments using the formula:

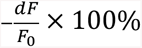

where *F_0_* refers to the QD photoluminescence at the initial temperature.

In solution, the PL intensity changed by 1.76 ± 0.21 % per °C, while for the surface-immobilized QDs, a slightly lower temperature sensitivity of 1.68 ± 0.01 % per °C was found. For later experiments with QD-coated substrates, the QD sensitivity determined on 2D surfaces was used.

### 3.4 Calibration of GNRs-induced thermoplasmonic heating

Before performing cell activity modulation experiments, the GNR-mediated plasmonic heating efficiency was calibrated using the CdSe/CdS@SiO_2_ QDs as local temperature probes. The GNR-QD-modified substrates were illuminated with a 785nm NIR laser (spot diameter of 230 µm – Figure 4g). A reference experiment was also performed on a QD-coated, GNR-free sample to check the effect of NIR illumination in the absence of the gold nanorods. The fluorescence intensity of the QDs was recorded for NIR laser power densities of 0.54 W/mm^2^, 2.78 W/mm^2^, and 5.11 W/mm^2^.

**Figure 4.**
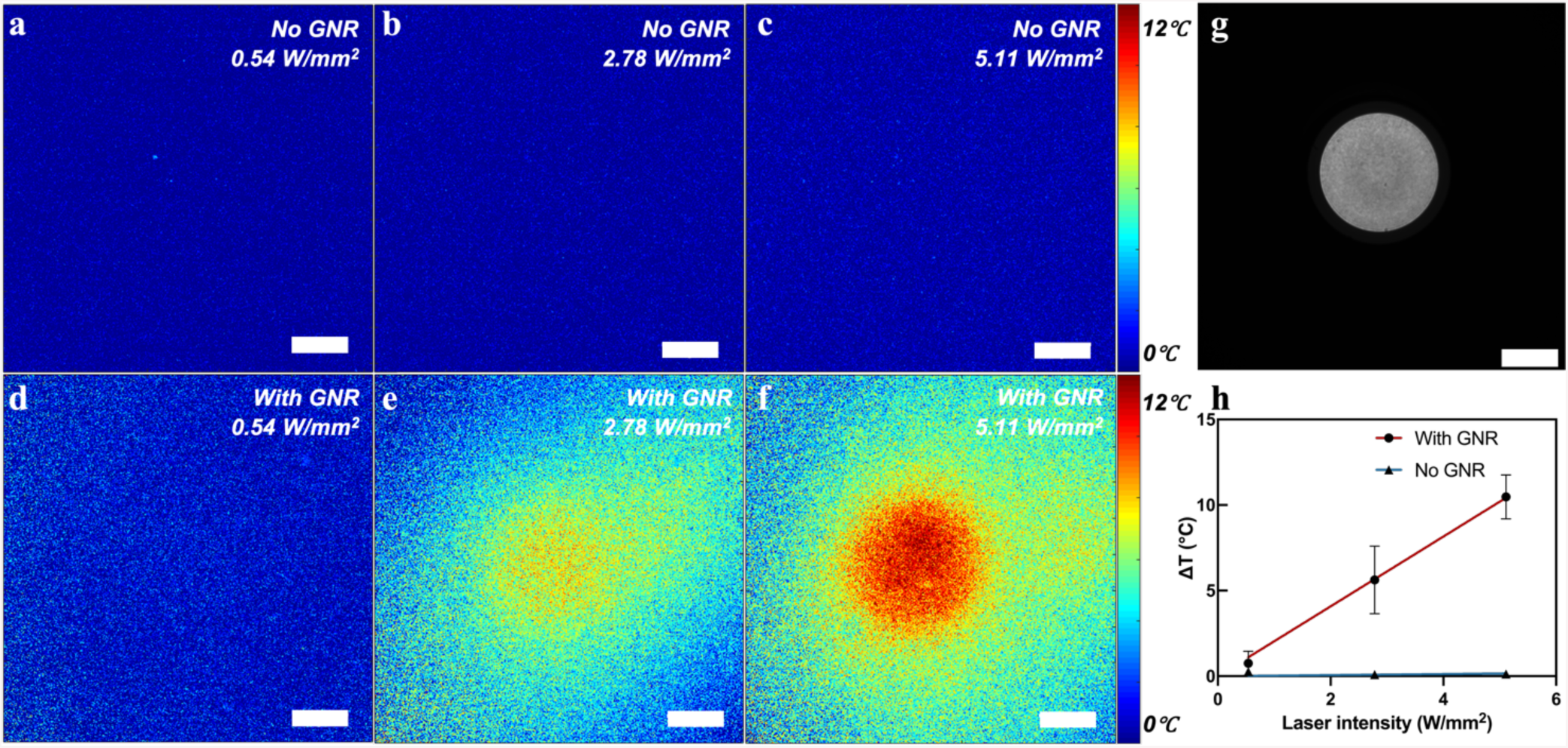
GNR plasmonic heating calibration. **a-c.** QD PL intensity maps of a GNR-free sample coated with QDs and illuminated with a 785 nm laser at intensities of (a) 0.54, (b) 2.78, (c) 5.11 W/mm2; **d-f.** 785nm light-induced plasmonic heating map of a GNR-QD-coated substrate for laser powers of 0.54, 2.78, and 5.11 W/mm2, respectively for panels d, e, and f. **g.** 785 nm light distribution. **h.** Temperature change inside the illuminated area as a function of the laser power (3 spots analyzed per laser intensity). Scale bar: 100 µm.

Figures 4a-f display the heat maps of the QD-modified sample (a-c) and GNR-QD-modified sample (d-f) obtained by subtracting the non-illuminated fluorescence image from the illuminated one in the same sample position as described in the experimental section. No distinct temperature changes were detected on GNR-free samples, while for the GNR-containing samples, the temperature in the illuminated area rose linearly with the NIR laser power, as shown in Figure 4h. A temperature change as high as 12 °C could be obtained for a NIR power density of 5.11 W/mm^2^. To monitor temperature evolution upon thermoplasmonic heating in function of time, QD photoluminescence was recorded under constant 2.78 W/mm^2^ NIR illumination power for 30 seconds. Under these conditions, Figure S5 (Supplementary Information) shows that the temperature increases within seconds upon NIR illumination to a steady state value.

### 3.5 Plasmonic modulation of HL-1 cell activities

Prior to performing plasmonic modulation experiments on HL-1 cell cultures, we observed cell activity upon NIR illumination on clean glass substrates in the absence of GNR, which demonstrated that NIR illumination on its own had no effect on cellular activity (results shown in Figure S6, Supporting Information). To investigate changes in cardiomyocyte activity by GNR-mediated plasmonic heating, we then cultured HL-1 cardiac muscle cells on GNR-QD modified substrates and recorded their beating activity by calcium imaging. Considering the activity differences observed in cultures with different cell densities (**Section 2.2**), we performed the modulation experiments on cultures with three different cell densities, i.e., single cells, small clusters, and confluent cell samples, respectively. Single-cell samples seeded initially with 10000/cm^2^ cells, presented mostly isolated cells and very few connected cells, with the total cell area being smaller than the NIR laser spot area. Small cluster samples seeded initially with 50000/cm^2^ cells, contained cell clusters covering an area slightly larger than that of the NIR laser spot, while confluent samples seeded initially with 100000/cm^2^ cells presented fully confluent cell layers over the whole area of the substrate. All the modulation experiments were performed on 2 DIV cultures (detailed protocol described in Experimental Section).

Figures 5a-c show representative fluorescence images of HL-1 cells loaded with Fluo-3 AM calcium dye for three different cell densities; each sample was illuminated with three different laser powers, 0.54 W/mm^2^, 2.78 W/mm^2^, and 5.11 W/mm^2^, corresponding to respectively 0.75 ± 0.41 °C, 5.63 ± 1.97 °C and 10.48 ± 1.29 °C local temperature increases at the sample surface according to the calibration described above. During the NIR modulation experiments, cells were kept in 1 × 10^−2^ M, pH 7.4 HEPES buffer with the culture chamber temperature maintained at 30 °C. The NIR laser illumination started 30 s after the beginning of image acquisition and lasted for 30 s, while the fluorescence signal was recorded for a total of 90 s (5 Hz sampling frequency - Figure 5d-f). An obvious calcium transient amplitude change was observed during NIR illumination, with a lower amplitude being observed alongside the higher beating rate. This effect results to a large extent from the low sampling frequency during recording and, to a smaller extent, from the temperature-dependent fluorescence of the calcium-sensitive dye.^46^ When the beating rate increases, low sampling frequency recordings can accurately convey the signal frequency but cannot reveal the full waveform of the calcium oscillation. The fluorescence traces were then FFT treated to convert time domain data into the frequency domain shown in Figure 5g-i. As shown in Figure 5, cultures with different cell densities responded differently to temperature changes. The changes in the beating rate of isolated cells correlated well with the applied NIR laser intensity, a higher laser intensity leading to faster beating. These cells displayed initially the slowest beating rate and with a NIR power density of 5.11 W/mm^2^ their activity became as fast as those of cells in confluent cultures.

**Figure 5.**
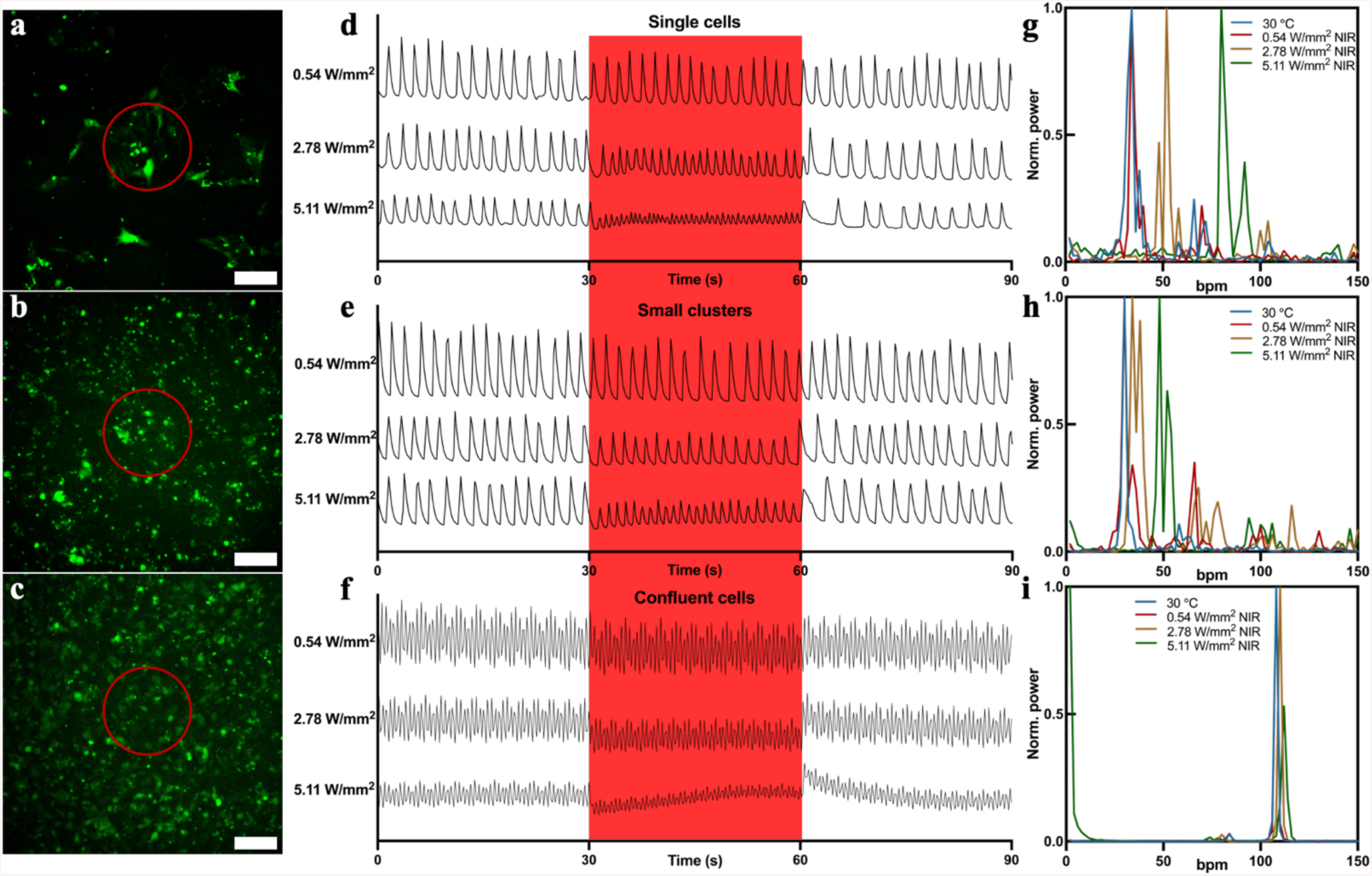
Calcium transients under thermoplasmonic modulation of HL-1 cultures. **a-c.** Fluorescence images of Fluo 3-loaded HL-1 cells (2 DIV) with different densities,(a) single cells, (b) small clusters, and (c) confluent cells. **d-f.** Calcium transients of cells in red circles before, during (red window) and after NIR irradiation. 3 samples per condition, with 3 measured areas per sample. **g-i.** Fast Fourier transform (FFT) analysis of beating rates. NIR-illuminated area is marked with a red circle. Scale bar: 100 µm.

When cell connectivity increased, the thermo-plasmonic effects became less pronounced. For small clusters, the initial beating rate at 30°C was similar to that observed for single cells, and also increased with NIR stimulation, but the beating rate changes were smaller than those observed for single cells. Given the intercellular connectivity, the cells outside the stimulated area influenced the beating rate of their heated neighbors. For confluent cell cultures, the beating rate modulation by plasmonic stimulation was even more limited, which may again be related to the influence of connected cells present outside of the stimulated area. The NIR illumination effects were reversible, with all cultures recovering their initial beating rate after stopping the illumination. Samples stimulated with high NIR powers displayed ‘overdrive suppression’ right after stimulation,^47,48^ which refers to a slower activity transient after stimulation at a rate faster than the intrinsic frequency before returning to the initial rate (visible from the last 10 seconds of the recorded signals). This behavior illustrates the power of the method to induce and study overdrive suppression mechanisms in vitro, relevant for arrhythmia treatment research.

The average evolution of HL-1 beating activity in response to plasmonic modulation is shown in Figures 6a-c (3 samples per condition, with 3 measured areas per sample), and the change in the beating rates for different NIR laser intensities is plotted in Figure 6d. Of all the three cell densities investigated, single cells are the most responsive to thermoplasmonic stimulation regardless of the NIR laser power, and the confluent cells are the least responsive. The trend seems opposite to the results discussed in section 2.1 for experiments performed with global temperature control, in which confluent cells responded more to increases in temperature. However, a careful analysis reveals that temperature effects are similar for NIR stimulated isolated cells in low density cultures. The 2.78 and 5.11 W/mm^2^ NIR illumination generate temperature changes of 5.63 ± 1.97 °C and 10.48 ± 1.29 °C, respectively, in the illuminated area. These correspond to extracellular temperature values of 35.63 °C (2.78 W/mm^2^ NIR) and 40.48 °C (5.11 W/mm^2^ NIR), respectively. In low density cultures, the beating rate of isolated cells increased with 7.6 bpm from 30 °C to 35 °C and 22 bpm from 30 °C to 40 °C in the global heating experiments (see Figure 2a), which were about the same as the bpm changes induced by the NIR illumination, i.e., 6.8 bpm change for 2.78 W/mm^2^ and 16 bpm change for 5.11 W/mm^2^, thus only slightly lower than during global heating. This small difference could be attributed to the heating localized on the substrate surface in NIR stimulation, meaning that the temperature change across the entire cell was slightly lower than in global heating experiments. On the other hand, for confluent cells, the plasmonic stimulation had a much lower effect than the global heating. This might be explained by the large constraint imposed by the many unstimulated cells in the synchronized community, as the beating behavior of cardiomyocytes follows the dominant rule of synchronization, meaning that local changes in beating rates are leveled out by the community behavior (i.e., the local temperature effects are canceled out by the global activity of the network).^49,50^ In future studies, it would be interesting to investigate the effect of patterned light stimulation on network synchronization patterns.^50^ To demonstrate the feasibility of this system in physiological conditions, we also conducted NIR modulation on HL-1 cultures with moderate NIR power (2.78 W/mm^2^ NIR) under 37 °C global culture temperature (Figure 6e). Similar to the results at 30 °C global temperature, single cells and small clusters respond more to the NIR modulation, while the confluent cells are the least responsive. Figure 6f shows the beating rate variation comparison between 37 °C and 30 °C global temperature both with 2.78 W/mm^2^ NIR stimulation, which indicates that single cells and small clusters respond more at 37 °C than at 30 °C global temperature, while confluent cells show the lowest activity changes to NIR stimulation regardless of the culture temperature.

**Figure 6.**
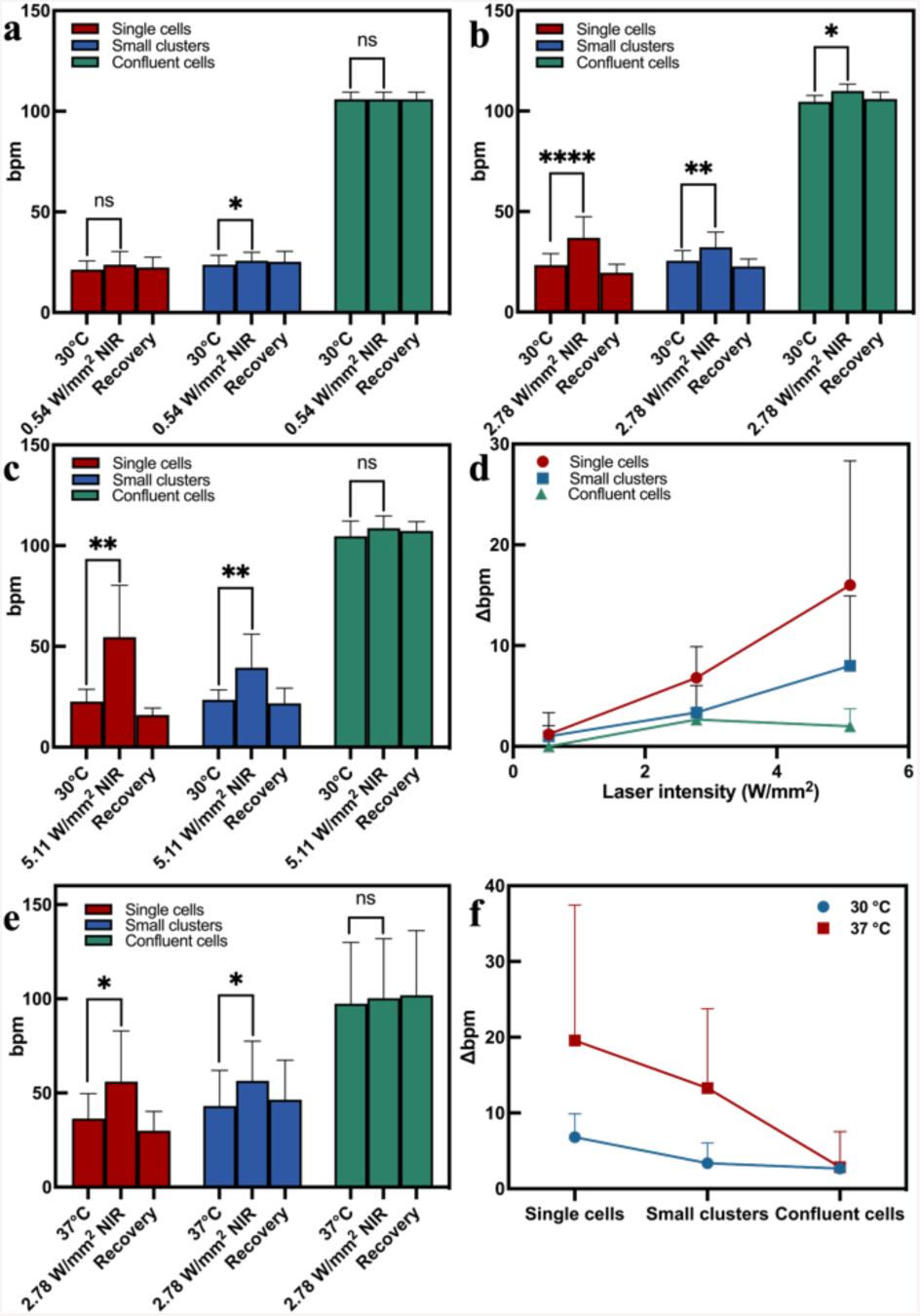
**a-c.** HL-1 beating rates for different culture densities at 30 °C global temperature under different NIR power densities: (a) 0.54 W/mm^2^, (b) 2.78 W/mm^2^ and (c) 5.11 W/mm^2^. p-values calculated from paired t-test, (*) p < 0.05, (**) p < 0.01, (***) p < 0.001, (****) p < 0.0001, (ns) not significant. **d.** Beating rate variation for three different cell densities as function of NIR laser power. Two-way ANOVA test: Cell density, p < 0.01, NIR laser power, p < 0.01. **e.** HL-1 beating rates for different culture densities at 37 °C global temperature under 2.78 W/mm^2^ NIR power. f. Beating rate variation under 2.78 W/mm^2^ NIR power at different global temperatures. Two-way ANOVA test: Temperature, p = 0.0138; Cell density, p = 0.0286.

### 3.6 Plasmon-controlled pacing depends on cell connectivity

Gap-junction connected cardiomyocytes display synchronous beating. In these experiments, we investigated whether the thermoplasmonic stimulated cells could act as a pacemaker to control the activity of connected cells outside the illuminated area. Therefore, we analyzed the beating activities of cells inside and outside the NIR illuminated area for samples with low and medium cell densities. The FFT analysis is shown in Figure 7. Cells in the areas delimited by the red circles were exposed directly to NIR light, while the cells in the areas delimited by the blue circles were not illuminated. For the small cluster sample, the beating activities observed for cells in these two regions changed in an identical way, indicating that in samples with gap-junction connected cells, the NIR illumination could be used to control the activity of cells outside the laser spot. For single-cell samples, where cells were not connected (as shown by their independent firing patterns – see Figure 7a), the plasmonic stimulation effects were highly localized. To better visualize the gap-junction between cells, fluorescence images of cell cultures after immunostaining of the gap-junction protein Connexin 43 were acquired for these two types of samples, see Figure S7, illustrating the gap-junction formation between cells also in small cluster samples.

**Figure 7.**
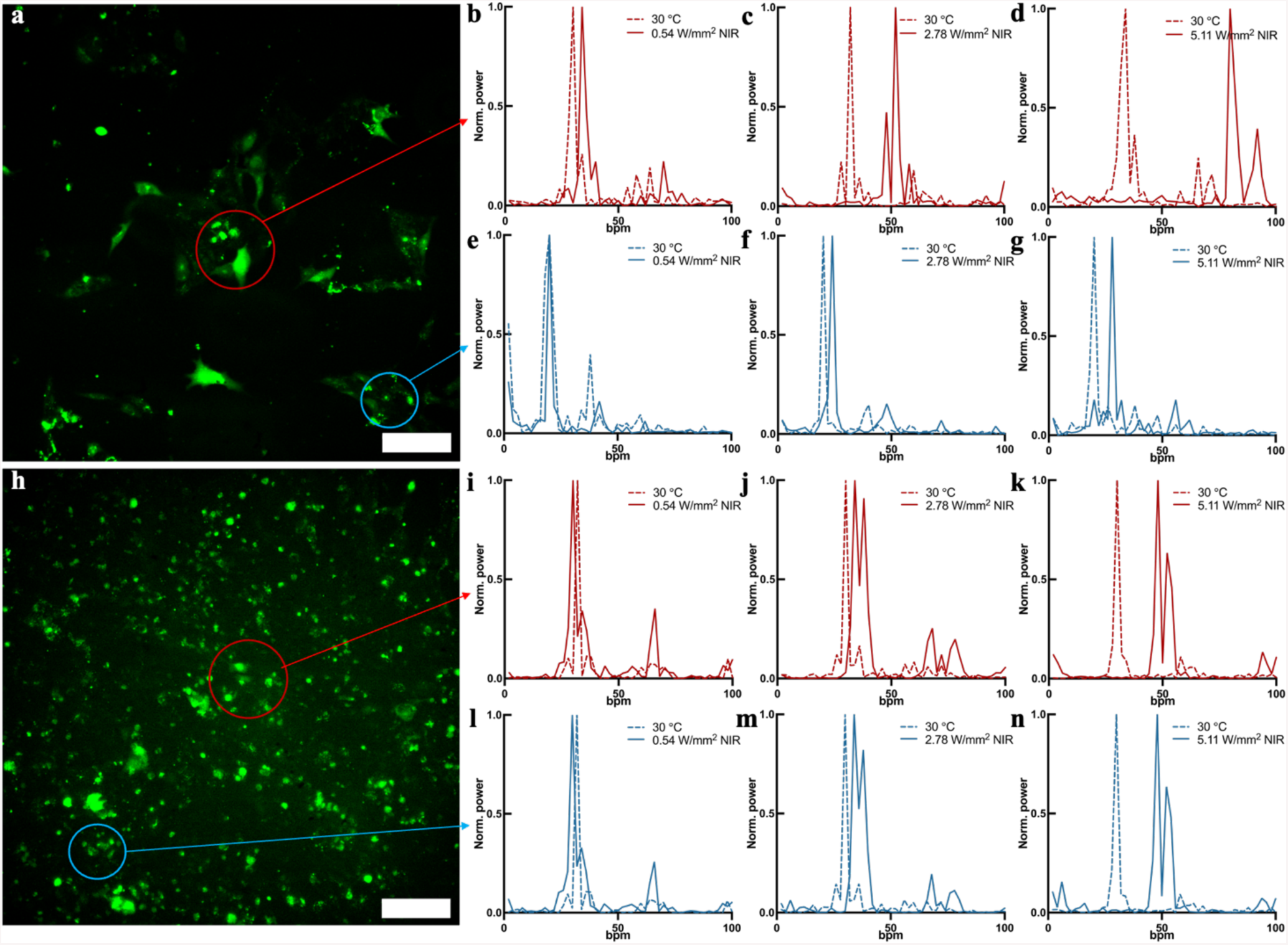
HL-1 cells synchronization upon thermoplasmonic stimulation. **a & h**. Fluorescence images of Fluo-3 AM-loaded HL-1 cells with (a) low connectivity (single-cell samples) and (h) moderate (small cluster samples) connectivity. **b-g**. Single-cell sample beating rate FFT analysis for cells inside the illuminated area (b-d), and outside the illuminated area (e-g). **i-n.** Small cluster sample beating rate FFT analysis for cells inside the illuminated area (i-k), and outside the illuminated area (l-n). Scale bar: 100 µm.

These results illustrate the potential of NIR-controlled plasmonic pacing in cardiomyocyte cultures and its relevance for arrhythmia and connectivity in vitro assays. The combination of plasmonic particles and temperature nanosensors allowed studying the link between the local temperature and cell activity.

## 4. CONCLUSIONS

In this paper, we discuss an all-optical system allowing cardiomyocyte activity modulation by NIR-controlled plasmon stimulation together with in situ local temperature monitoring. Cell beating behavior was modulated through GNR-mediated photothermal effect, while the local extracellular temperature was monitored through variations in the photoluminescence of QDs. CdSe/CdS@SiO_2_ QDs demonstrate good photostability and temperature sensitivity of about 1.68% per °C under physiological conditions. The amount of heat generated by GNRs could be measured as a function of NIR laser power and linked to changes in cardiomyocyte beating rates. QD temperature-modulated fluorescence was used to generate substrate temperature maps, revealing that local temperature changes of up to 10 °C were produced by NIR-plasmonic heating at moderate power levels (5.11 W/mm^2^). Experiments on HL-1 cardiac cells showed that cells changed their beating rates in response to plasmonic heating and that the temperature effects depended largely on cell connectivity. Remarkably, in the presence of the QD temperature probes, the local temperature could be monitored with high spatial and thermal resolution, allowing controlling plasmonic heating during modulation experiments. This all-optical, nanoparticle-mediated, stimulation system also opens the possibility to pace cardiomyocyte activity via plasmonic heating in arrythmia studies and is also translatable to 3D cell and tissue constructs.

## ASSOCIATED CONTENT

### Supporting Information

The Supporting Information is available free of charge.

Supporting figures including AFM and TEM images, UV-Vis spectra, fluorescence images, etc. (DOC)

## AUTHOR INFORMATION

### Notes

The authors declare no conflict of interest.

## ACKNOWLEDGEMENTS

The authors thank prof. dr. Zeger Hens and dr. Tangi Aubert for kindly supplying the silica-coated CdSe/CdS quantum dots. F. Guo acknowledges the financial support from the China Scholarship Council (CSC). S.Jooken acknowledges financial support from the Research Foundation - Flanders, Belgium (Grant no.1SC3819N). L. Liu acknowledges financial support from the Research Foundation - Flanders, Belgium (Grant no.12V4422N). C. Bartic and O. Deschaume acknowledge the financial support from the Research Foundation - Flanders, Belgium (Grant no. G0947.17N). W. Thielemans, C. Bartic and O. Deschaume acknowledge the financial support from the Onderzoeksraad, KU Leuven, Belgium (Grant no. C14/18/061).

## REFERENCES

1. Wei, F.; Pourrier, M.; Strauss, D. G.; Stockbridge, N.; Pang, L. Effects of Electrical Stimulation on HiPSC-CM Responses to Classic Ion Channel Blockers. Toxicological Sciences 2020, 174 (2), 254–265. 10.1093/toxsci/kfaa010.

2. Stoppel, W. L.; Kaplan, D. L.; Black, L. D. Electrical and Mechanical Stimulation of Cardiac Cells and Tissue Constructs. Adv Drug Deliv Rev 2016, 96, 135–155. 10.1016/j.addr.2015.07.009.

3. Tandon, N.; Cannizzaro, C.; Chao, P.-H. G.; Maidhof, R.; Marsano, A.; Au, H. T. H.; Radisic, M.; Vunjak-Novakovic, G. Electrical Stimulation Systems for Cardiac Tissue Engineering. Nat Protoc 2009, 4 (2), 155–173. 10.1038/nprot.2008.183.

4. Hirt, M. N.; Boeddinghaus, J.; Mitchell, A.; Schaaf, S.; Börnchen, C.; Müller, C.; Schulz, H.; Hubner, N.; Stenzig, J.; Stoehr, A.; Neuber, C.; Eder, A.; Luther, P. K.; Hansen, A.; Eschenhagen, T. Functional Improvement and Maturation of Rat and Human Engineered Heart Tissue by Chronic Electrical Stimulation. J Mol Cell Cardiol 2014, 74, 151–161. 10.1016/j.yjmcc.2014.05.009.

5. Jia, Z.; Valiunas, V.; Lu, Z.; Bien, H.; Liu, H.; Wang, H.-Z.; Rosati, B.; Brink, P. R.; Cohen, I. S.; Entcheva, E. Stimulating Cardiac Muscle by Light: Cardiac Optogenetics by Cell Delivery. Circ Arrhythm Electrophysiol 2011, 4 (5), 753–760. 10.1161/CIRCEP.111.964247.

6. Bruegmann, T.; Malan, D.; Hesse, M.; Beiert, T.; Fuegemann, C.; Fleischmann, B.; Sasse, P. Optogenetic Control of Heart Muscle in Vitro and in Vivo. Nature methods 2010, 7, 897–900. 10.1038/nmeth.1512.

7. Entcheva, E. Cardiac Optogenetics. American Journal of Physiology-Heart and Circulatory Physiology 2013, 304 (9), H1179–H1191. 10.1152/ajpheart.00432.2012.

8. Ambrosi, C. M.; Klimas, A.; Yu, J.; Entcheva, E. Cardiac Applications of Optogenetics. Prog Biophys Mol Biol 2014, 115 (2–3), 294–304. 10.1016/j.pbiomolbio.2014.07.001.

9. Williams, J. C.; Entcheva, E. Optogenetic versus Electrical Stimulation of Human Cardiomyocytes: Modeling Insights. Biophys J 2015, 108 (8), 1934–1945. 10.1016/j.bpj.2015.03.032.

10. Floria, M.; Radu, S.; Gosav, E. M.; Moraru, A. C.; Serban, T.; Carauleanu, A.; Costea, C. F.; Ouatu, A.; Ciocoiu, M.; Tanase, D. M. Cardiac Optogenetics in Atrial Fibrillation: Current Challenges and Future Opportunities. BioMed Research International 2020, 2020, e8814092. 10.1155/2020/8814092.

11. Riefolo, F.; Matera, C.; Garrido-Charles, A.; Gomila, A. M. J.; Sortino, R.; Agnetta, L.; Claro, E.; Masgrau, R.; Holzgrabe, U.; Batlle, M.; Decker, M.; Guasch, E.; Gorostiza, P. Optical Control of Cardiac Function with a Photoswitchable Muscarinic Agonist. J. Am. Chem. Soc. 2019, 141 (18), 7628– 7636. 10.1021/jacs.9b03505.

12. Duran-Corbera, A.; Faria, M.; Ma, Y.; Prats, E.; Dias, A.; Catena, J.; Martinez, K. L.; Raldua, D.; Llebaria, A.; Rovira, X. A Photoswitchable Ligand Targeting the Β1-Adrenoceptor Enables Light-Control of the Cardiac Rhythm**. Angewandte Chemie International Edition 2022, 61 (30), e202203449. 10.1002/anie.202203449.

13. Parameswaran, R.; Koehler, K.; Rotenberg, M. Y.; Burke, M. J.; Kim, J.; Jeong, K.-Y.; Hissa, B.; Paul, M. D.; Moreno, K.; Sarma, N.; Hayes, T.; Sudzilovsky, E.; Park, H.-G.; Tian, B. Optical Stimulation of Cardiac Cells with a Polymer-Supported Silicon Nanowire Matrix. Proceedings of the National Academy of Sciences 2019, 116 (2), 413–421. 10.1073/pnas.1816428115.

14. Lodola, F.; Vurro, V.; Crasto, S.; Di Pasquale, E.; Lanzani, G. Optical Pacing of Human-Induced Pluripotent Stem Cell-Derived Cardiomyocytes Mediated by a Conjugated Polymer Interface. Advanced Healthcare Materials 2019, 8 (13), 1900198. 10.1002/adhm.201900198.

15. Savchenko, A.; Cherkas, V.; Liu, C.; Braun, G. B.; Kleschevnikov, A.; Miller, Y. I.; Molokanova, E. Graphene Biointerfaces for Optical Stimulation of Cells. Science Advances 2018, 4 (5), eaat0351. 10.1126/sciadv.aat0351.

16. Bruno, G.; Melle, G.; Barbaglia, A.; Iachetta, G.; Melikov, R.; Perrone, M.; Dipalo, M.; De Angelis, F. All-Optical and Label-Free Stimulation of Action Potentials in Neurons and Cardiomyocytes by Plasmonic Porous Metamaterials. Advanced Science 2021, 8 (21), 2100627. 10.1002/advs.202100627.

17. Marino, A.; Arai, S.; Hou, Y.; Degl’Innocenti, A.; Cappello, V.; Mazzolai, B.; Chang, Y.-T.; Mattoli, V.; Suzuki, M.; Ciofani, G. Gold Nanoshell-Mediated Remote Myotube Activation. ACS Nano 2017, 11 (3), 2494–2508. 10.1021/acsnano.6b08202.

18. Gentemann, L.; Kalies, S.; Coffee, M.; Meyer, H.; Ripken, T.; Heisterkamp, A.; Zweigerdt, R.; Heinemann, D. Modulation of Cardiomyocyte Activity Using Pulsed Laser Irradiated Gold Nanoparticles. *Biomed. Opt. Express*, BOE 2017, 8 (1), 177–192. 10.1364/BOE.8.000177.

19. Fang, J.; Liu, D.; Xu, D.; Wu, Q.; Li, H.; Li, Y.; Hu, N. Integrated Au-Nanoroded Biosensing and Regulating Platform for Photothermal Therapy of Bradyarrhythmia. Research 2022, 2022. 10.34133/2022/9854342.

20. Smith, N. I.; Kumamoto, Y.; Iwanaga, S.; Ando, J.; Fujita, K.; Kawata, S. A Femtosecond Laser Pacemaker for Heart Muscle Cells. *Opt. Express*, OE 2008, 16 (12), 8604–8616. 10.1364/OE.16.008604.

21. Yeh, Y.-C.; Creran, B.; Rotello, V. M. Gold Nanoparticles: Preparation, Properties, and Applications in Bionanotechnology. Nanoscale 2012, 4 (6), 1871–1880. 10.1039/C1NR11188D.

22. Cao, J.; Sun, T.; Grattan, K. T. V. Gold Nanorod-Based Localized Surface Plasmon Resonance Biosensors: A Review. Sensors and Actuators B: Chemical 2014, 195, 332–351. 10.1016/j.snb.2014.01.056.

23. Bendix, P. M.; Reihani, S. N. S.; Oddershede, L. B. Direct Measurements of Heating by Electromagnetically Trapped Gold Nanoparticles on Supported Lipid Bilayers. ACS Nano 2010, 4 (4), 2256–2262. 10.1021/nn901751w.

24. Jang, H.; Yoon, D.; Nam, Y. Enhancement of Thermoplasmonic Neural Modulation Using a Gold Nanorod-Immobilized Polydopamine Film. ACS Appl. Mater. Interfaces 2022, 14 (21), 24122–24132. 10.1021/acsami.2c03289.

25. Yoo, S.; Hong, S.; Choi, Y.; Park, J.-H.; Nam, Y. Photothermal Inhibition of Neural Activity with Near-Infrared-Sensitive Nanotransducers. ACS Nano 2014, 8 (8), 8040–8049. 10.1021/nn5020775.

26. Biju, V.; Makita, Y.; Sonoda, A.; Yokoyama, H.; Baba, Y.; Ishikawa, M. Temperature-Sensitive Photoluminescence of CdSe Quantum Dot Clusters. J. Phys. Chem. B 2005, 109 (29), 13899–13905. 10.1021/jp050424l.

27. Dai, Q.; Song, Y.; Li, D.; Chen, H.; Kan, S.; Zou, B.; Wang, Y.; Deng, Y.; Hou, Y.; Yu, S.; Chen, L.; Liu, B.; Zou, G. Temperature Dependence of Band Gap in CdSe Nanocrystals. Chemical Physics Letters 2007, 439 (1), 65–68. 10.1016/j.cplett.2007.03.034.

28. Maestro, L. M.; Rodríguez, E. M.; Rodríguez, F. S.; la Cruz, M. C. I.; Juarranz, A.; Naccache, R.; Vetrone, F.; Jaque, D.; Capobianco, J. A.; Solé, J. G. CdSe Quantum Dots for Two-Photon Fluorescence Thermal Imaging. Nano Lett. 2010, 10 (12), 5109–5115. 10.1021/nl1036098.

29. Gerion, D.; Pinaud, F.; Williams, S. C.; Parak, W. J.; Zanchet, D.; Weiss, S.; Alivisatos, A. P. Synthesis and Properties of Biocompatible Water-Soluble Silica-Coated CdSe/ZnS Semiconductor Quantum Dots. J. Phys. Chem. B 2001, 105 (37), 8861–8871. 10.1021/jp0105488.

30. Pérez-Juste, J.; Liz-Marzán, L. M.; Carnie, S.; Chan, D. Y. C.; Mulvaney, P. Electric-Field-Directed Growth of Gold Nanorods in Aqueous Surfactant Solutions. Advanced Functional Materials 2004, 14 (6), 571–579. 10.1002/adfm.200305068.

31. Jana, N. R.; Gearheart, L.; Murphy, C. J. Seeding Growth for Size Control of 5−40 Nm Diameter Gold Nanoparticles. Langmuir 2001, 17 (22), 6782–6786. 10.1021/la0104323.

32. Yu, W.; Deschaume, O.; Dedroog, L.; Garcia Abrego, C. J.; Zhang, P.; Wellens, J.; de Coene, Y.; Jooken, S.; Clays, K.; Thielemans, W.; Glorieux, C.; Bartic, C. Light-Addressable Nanocomposite Hydrogels Allow Plasmonic Actuation and In Situ Temperature Monitoring in 3D Cell Matrices. Advanced Functional Materials 2022, 32 (5), 2108234. 10.1002/adfm.202108234.

33. Mehtala, J. G.; Zemlyanov, D. Y.; Max, J. P.; Kadasala, N.; Zhao, S.; Wei, A. Citrate-Stabilized Gold Nanorods. Langmuir 2014, 30 (46), 13727–13730. 10.1021/la5029542.

34. Cirillo, M.; Aubert, T.; Gomes, R.; Van Deun, R.; Emplit, P.; Biermann, A.; Lange, H.; Thomsen, C.; Brainis, E.; Hens, Z. “Flash” Synthesis of CdSe/CdS Core–Shell Quantum Dots. Chem. Mater. 2014, 26 (2), 1154–1160. 10.1021/cm403518a.

35. Aubert, T.; Soenen, S. J.; Wassmuth, D.; Cirillo, M.; Van Deun, R.; Braeckmans, K.; Hens, Z. Bright and Stable CdSe/CdS@SiO2 Nanoparticles Suitable for Long-Term Cell Labeling. ACS Appl. Mater. Interfaces 2014, 6 (14), 11714–11723. 10.1021/am502367b.

36. Jooken, S.; Coene, Y. de; Deschaume, O.; Zámbó, D.; Aubert, T.; Hens, Z.; Dorfs, D.; Verbiest, T.; Clays, K.; Callewaert, G.; Bartic, C. Enhanced Electric Field Sensitivity of Quantum Dot/Rod Two-Photon Fluorescence and Its Relevance for Cell Transmembrane Voltage Imaging. Nanophotonics 2021, 10 (9), 2407–2420. 10.1515/nanoph-2021-0077.

37. Drijvers, E.; Liu, J.; Harizaj, A.; Wiesner, U.; Braeckmans, K.; Hens, Z.; Aubert, T. Efficient Endocytosis of Inorganic Nanoparticles with Zwitterionic Surface Functionalization. ACS Appl. Mater. Interfaces 2019, 11 (42), 38475–38482. 10.1021/acsami.9b12398.

38. Miranda, A.; Martínez, L.; De Beule, P. A. A. Facile Synthesis of an Aminopropylsilane Layer on Si/SiO2 Substrates Using Ethanol as APTES Solvent. MethodsX 2020, 7, 100931. 10.1016/j.mex.2020.100931.

39. White, S. M.; Constantin, P. E.; Claycomb, W. C. Cardiac Physiology at the Cellular Level: Use of Cultured HL-1 Cardiomyocytes for Studies of Cardiac Muscle Cell Structure and Function. American Journal of Physiology-Heart and Circulatory Physiology 2004, 286 (3), H823–H829. 10.1152/ajpheart.00986.2003.

40. Rueden, C. T.; Schindelin, J.; Hiner, M. C.; DeZonia, B. E.; Walter, A. E.; Arena, E. T.; Eliceiri, K. W. ImageJ2: ImageJ for the next Generation of Scientific Image Data. BMC Bioinformatics 2017, 18 (1), 529. 10.1186/s12859-017-1934-z.

41. Uchida, T.; Kitora, R.; Gohara, K. Temperature Dependence of Synchronized Beating of Cultured Neonatal Rat Heart-Cell Networks with Increasing Age Measured by Multi-Electrode Arrays. Trends Med 2018, 18 (4). 10.15761/TiM.1000145.

42. Chen, Y.-J.; Chen, Y.-C.; Chan, P.; Lin, C.-I.; Chen, S.-A. Temperature Regulates the Arrhythmogenic Activity of Pulmonary Vein Cardiomyocytes. J Biomed Sci 2003, 10 (5), 535–543. 10.1007/BF02256115.

43. Fu, Y.; Zhang, G.-Q.; Hao, X.-M.; Wu, C.-H.; Chai, Z.; Wang, S.-Q. Temperature Dependence and Thermodynamic Properties of Ca2+ Sparks in Rat Cardiomyocytes. Biophys J 2005, 89 (4), 2533–2541. 10.1529/biophysj.105.067074.

44. Oyamada, M.; Kimura, H.; Oyamada, Y.; Miyamoto, A.; Ohshika, H.; Mori, M. The Expression, Phosphorylation, and Localization of Connexin 43 and Gap-Junctional Intercellular Communication during the Establishment of a Synchronized Contraction of Cultured Neonatal Rat Cardiac Myocytes. Experimental Cell Research 1994, 212 (2), 351–358. 10.1006/excr.1994.1154.

45. Kojima, K.; Kaneko, T.; Yasuda, K. Role of the Community Effect of Cardiomyocyte in the Entrainment and Reestablishment of Stable Beating Rhythms. Biochemical and Biophysical Research Communications 2006, 351 (1), 209–215. 10.1016/j.bbrc.2006.10.037.

46. Oliver, A. E.; Baker, G. A.; Fugate, R. D.; Tablin, F.; Crowe, J. H. Effects of Temperature on Calcium-Sensitive Fluorescent Probes. Biophysical Journal 2000, 78 (4), 2116–2126. 10.1016/S0006-3495(00)76758-0.

47. Overdrive suppression of spontaneously beating chick heart cell aggregates: experiment and theory. 10.1152/ajpheart.1995.269.3.H1153.

48. Vassalle, M. The Relationship among Cardiac Pacemakers. Overdrive Suppression. Circ Res 1977, 41 (3), 269–277. 10.1161/01.RES.41.3.269.

49. Hayashi, T.; Tokihiro, T.; Kurihara, H.; Yasuda, K. Community Effect of Cardiomyocytes in Beating Rhythms Is Determined by Stable Cells. Sci Rep 2017, 7 (1), 15450. 10.1038/s41598-017-15727-5.

50. Yasuda, K. Dominant Rule of Community Effect in Synchronized Beating Behavior of Cardiomyocyte Networks. Biophys Rev 2020, 12 (2), 481–501. 10.1007/s12551-020-00688-3.

