## Supporting information for "Thermoplasmonic modulation of cardiomyocytes activity with local temperature read-out"

**Overview of supporting information**

**Figure S1 AFM and UV-visible spectrum of GNRs**

**Figure S2 TEM and Extinction/Emission spectra of CdSe/CdS@$SiO_2$ QDs**

**Figure S3 AFM of an APTES-GNR modified glass surface**

**Figure S4 Calibration of CdSe/CdS@$SiO_2$ QD temperature sensitivity**

**Figure S5 QD fluorescence variation upon plasmonic heating.**

**Figure S6 NIR stimulation on 2 DIV cultured HL-1 cells on clean glass**

**Figure S7 Immunofluorescence microscopy of gap-junction protein Connexin 43.**

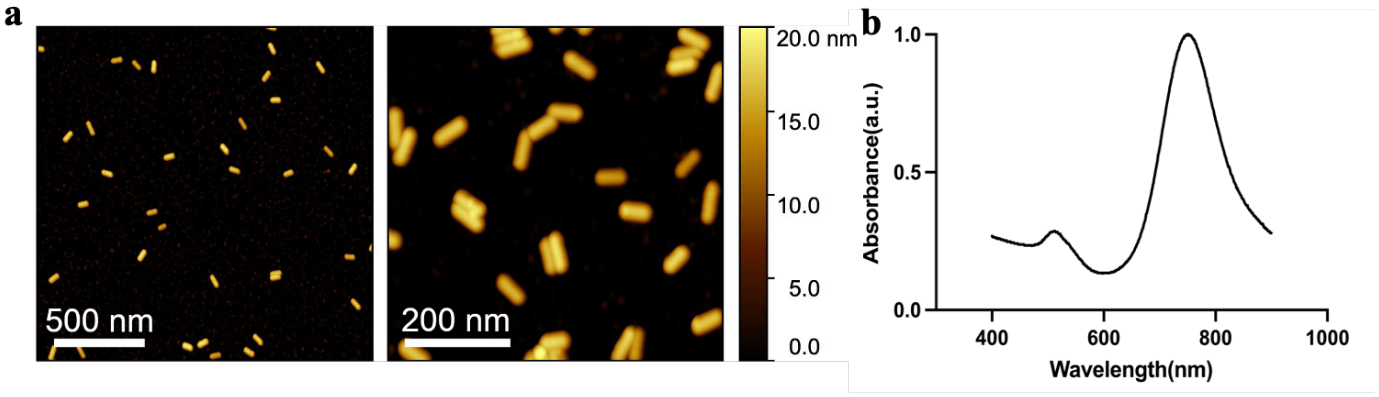


**Figure S1.** (a) AFM images and (b) UV-visible spectrum of GNRs.

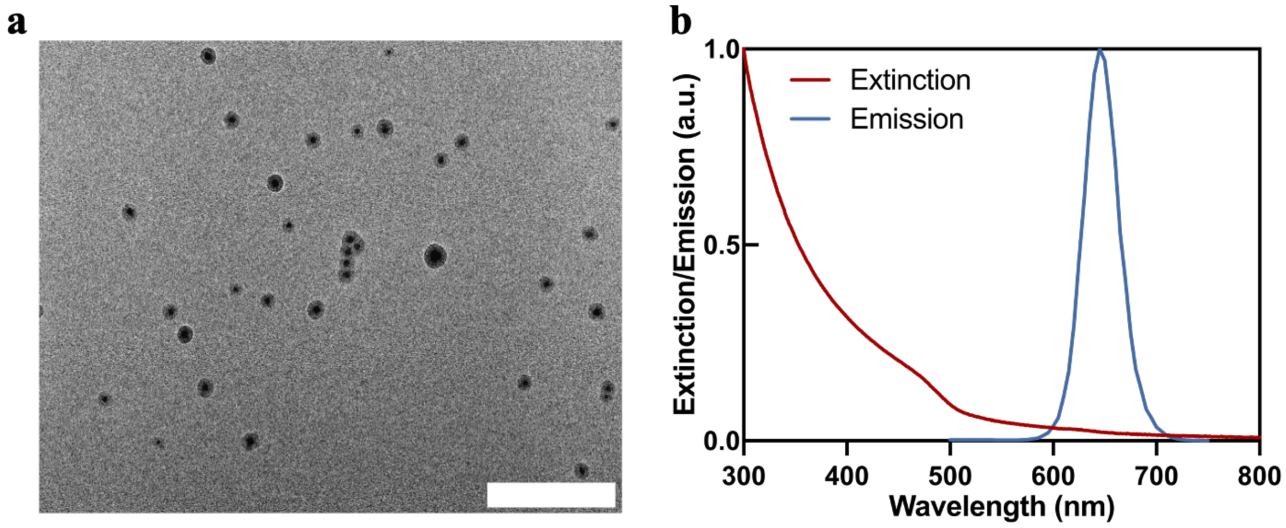


**Figure S2.** (a) TEM image - Scale bar: 200 nm and (b) Extinction/Emission spectra of CdSe/CdS@$SiO_2$ QDs.

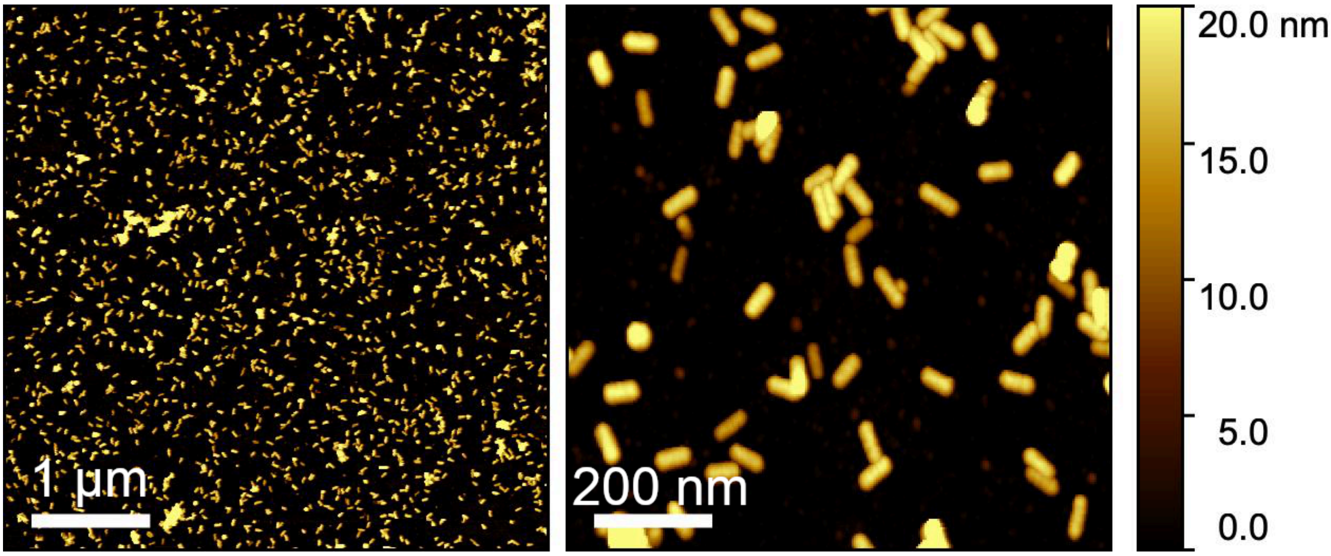


**Figure S3.** AFM of an APTES-GNR modified glass surface.

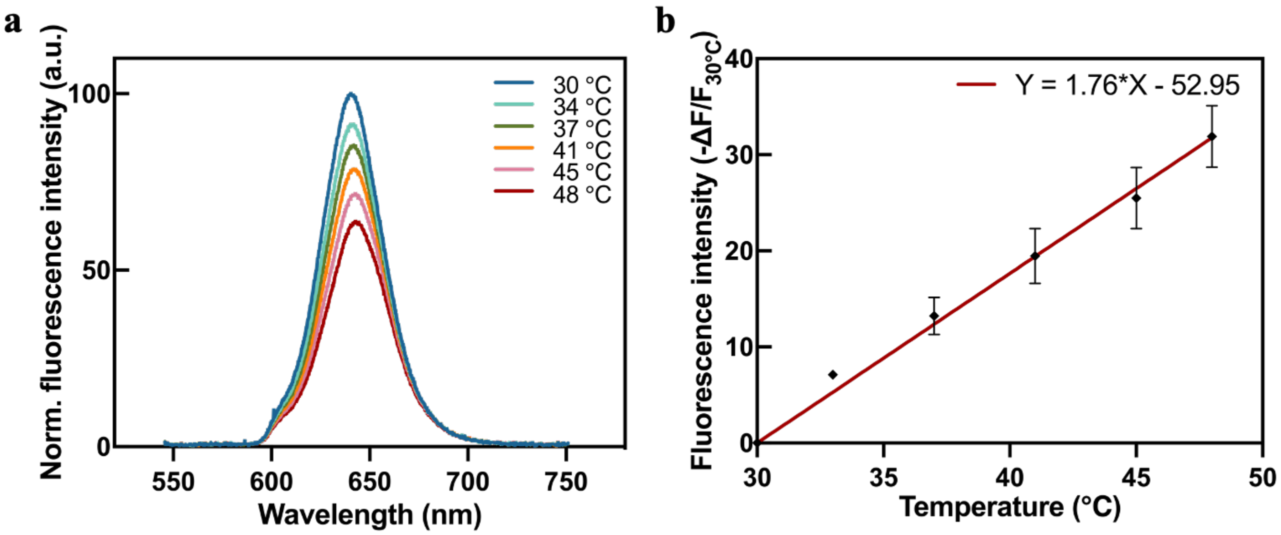


**Figure S4.** Calibration of CdSe/CdS@$SiO_2$ QD temperature sensitivity. **a.** Spectra of CdSe/CdS@$SiO_2$ dispersed in $1 \times 10^{-2}$ M, pH 7.4 HEPES buffer for temperatures between 30 and 48 °C. **b.** Percentual change of PL intensity of the CdSe/CdS@$SiO_2$ QD solution as a function of solution temperature, calculated from the data shown in panel a.

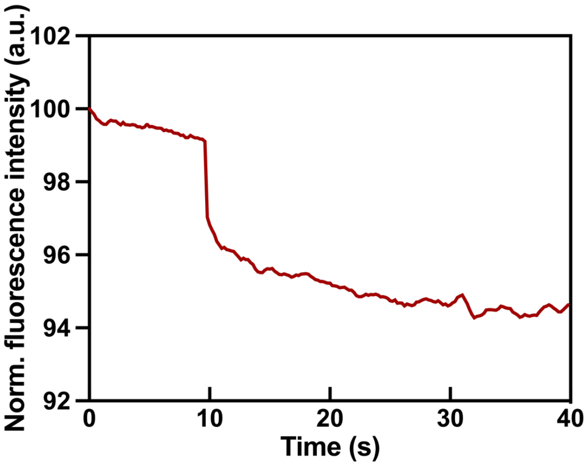


**Figure S5.** QD fluorescence intensity change upon NIR illumination as a function of time. NIR illumination starts from 10 seconds.

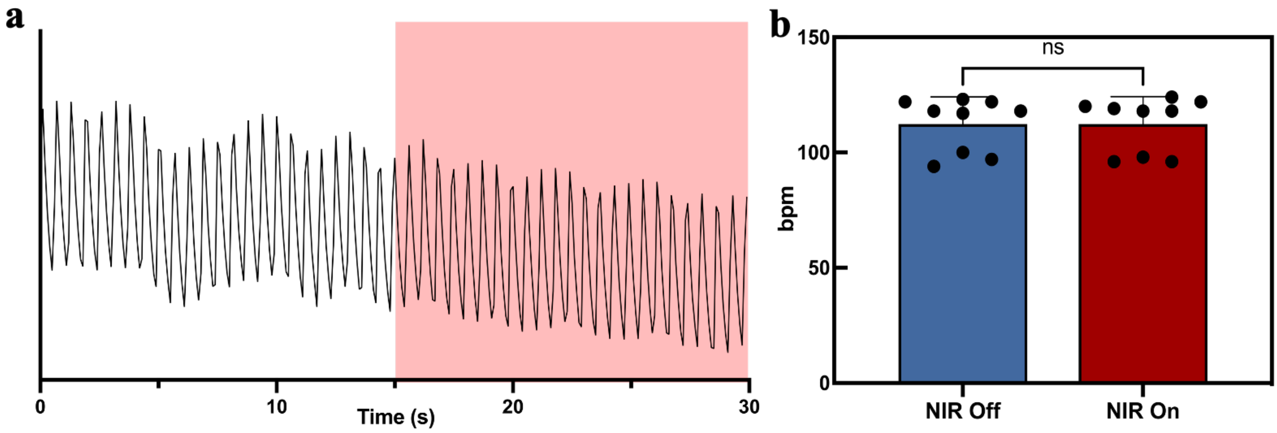


**Figure S6.** NIR stimulation on 2 DIV cultured HL-1 cells on clean glass, without GNR, environmental temperature 30 °C, NIR power 5.11W/mm$^2$. **a.** HL-1 calcium transients before and during (pink window) NIR irradiation. **b.** Statistical results for HL-1 cell beating rate variation with and without NIR illumination. p-value calculated from paired t-test, (ns) not significant, N = 9.

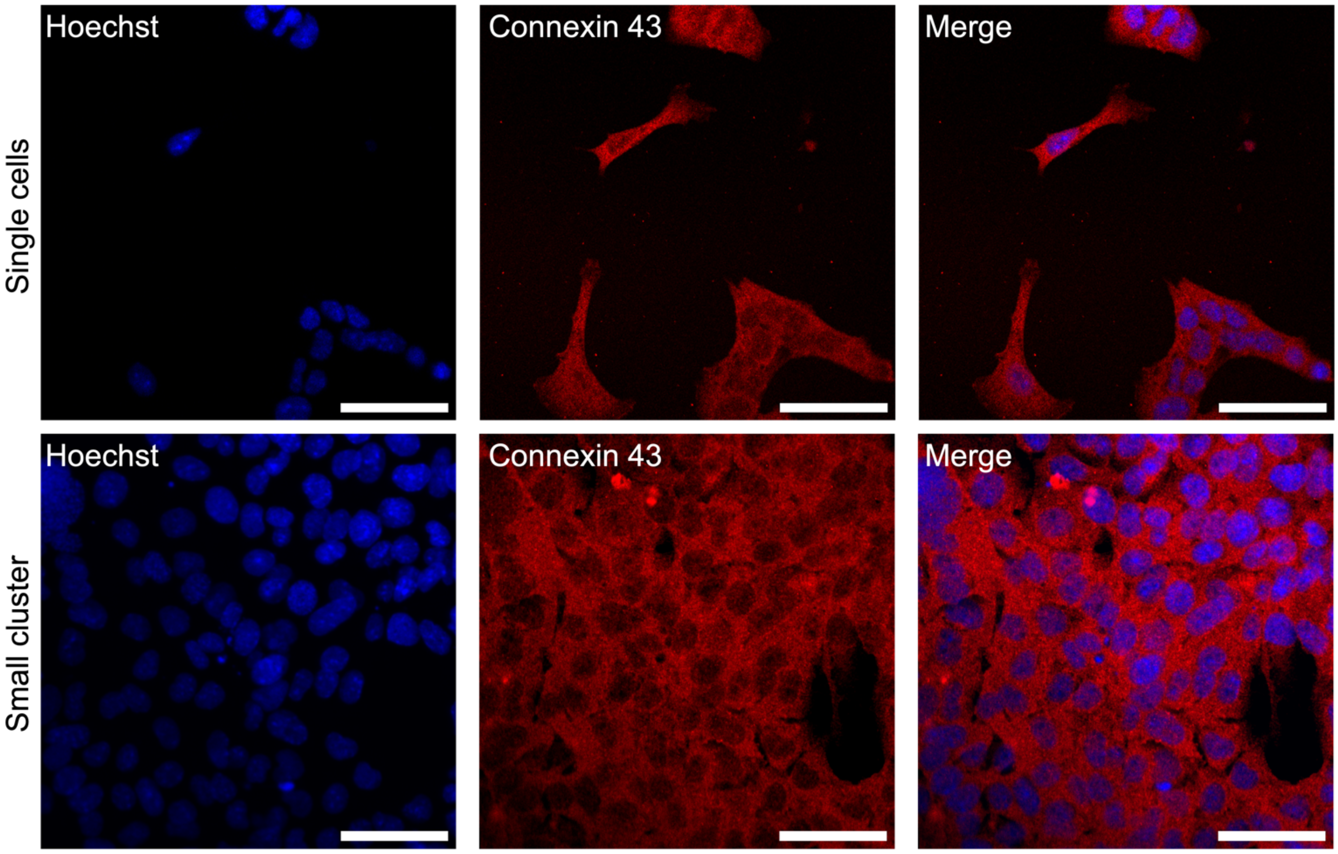


**Figure S7.** Immunofluorescence microscopy of gap-junction protein Connexin 43 on 2DIV HL-1 cell cultures. Scale bar: 50 µm.
